# High-throughput ligand profile characterization in novel biosensor cell lines expressing seven heterologous olfactory receptors for the detection of volatile plant biomarkers

**DOI:** 10.1101/2023.04.28.538535

**Authors:** Katalin Zboray, Ádám V. Tóth, Tímea D. Miskolczi, Emilio Casanova, Árpád Mike, László Sági, Péter Lukács

## Abstract

Agriculturally important crop plants emit a multitude of volatile organic compounds (VOCs), which are excellent indicators of their health status and/or their interaction with pathogens and pests. Here, we present the generation of a novel cellular biosensor panel for the recognition of fungal pathogen-related VOCs we had identified in the field as well as during controlled inoculations of several crop plants. The panel consists of seven stable HEK293 cell lines each expressing a functional *Drosophila* olfactory receptor as a biosensing element and a genetically encoded fluorescent calcium indicator protein. For high-throughput measurement of odorant binding by the cells, fluorescence response was detected in an automated 384-well microplate reader upon the injection of tester VOCs. Biosensor cell lines were characterized for their reference ligand binding in more detail, then a set of 66 VOCs was profiled on all cell lines over a concentration range of three orders of magnitude (1 μM to 100 μM). Forty-six VOCs (70%) evoked a response in at least one biosensor cell line and certain VOCs could activate the cell lines already from the nanomolar (ppb) concentration. Interaction profiles mapped in this study will support biosensor development for agricultural applications, but the olfactory receptor proteins may also be purified from these cell lines at sufficient yields for further processing including structure determination or coupling with artificial sensor devices.

## 1. Introduction

Volatile organic compound (VOC) composition of airborne emissions represents valuable information in countless scenarios from the detection of explosives or toxic material (Glatz and Bailey-Hill, 2011) to the distinction between healthy and diseased biological samples based on the presence or modification of a certain VOC blend (Wilson, 2016).

In the animal kingdom, VOCs are bound and recognized by odorant or olfactory receptors (OR), although several other peptide molecules such as Odorant Binding Proteins (OBP) are also capable of specific VOC recognition. It is due to these receptors that animals, such as drug-sniffing dogs, can detect and trace marker volatiles with unprecedented sensitivity. Mammals have several hundred to a thousand ORs, which are G-protein-coupled receptors (GPCRs) (Kato and Touhara, 2009). Numerous research programs have focused on the development of biosensors based on such mammalian receptors. However, due to their mechanism of action, the receptor itself is not sufficient for odorant perception, and additional signal transduction molecules are required for their proper function (Peterlin et al., 2014).

Insect ORs have a substantially different structure and mode of operation. For insects, too, odor perception is critical for feeding, oviposition, mate recognition, and predator avoidance (Carey and Carlson, 2011); therefore, insects are also able to sense relevant odorants at minute concentrations. Moreover, they possess a much narrower receptor repertoire compared to mammals, and – unlike GPCRs – do not need additional components for odorant recognition (Liu et al., 2014). Insect ORs are typically assembled by two types of proteins: the tuning OR, which is responsible for VOC binding and ligand-specificity, and the Olfactory Receptor Co-receptor (ORCO, previously OR83b; Vosshall and Hansson, 2011) that is essential for proper OR folding and function. While the amino acid sequence of OR proteins is very diverse, that of ORCO proteins is highly conserved even among diverse species. In many cases OR and ORCO protein combinations from different species can form functional channels. Cryo-EM structure determination of the ORCO protein from a parasitic fig wasp (*Apocrypta bakeri*) revealed a heterotetrameric structure in the resulting autonomous cation channel (Butterwick et al., 2018). Regarding their mechanism of action insect ORs predominantly work as ligand-gated ion channels (Sato et al., 2008; Wicher et al., 2008), although G-protein-mediated signaling was also indicated in several studies (Getahun et al., 2013; Nakagawa and Vosshall, 2009; Neuhaus et al., 2011; Sato et al., 2008; Silbering and Benton, 2010). VOC ligand-binding induces channel opening and an influx of cations. Channel permeability is much higher for monovalent (e.g., Na^+^, K^+,^ and Cs^+^) than for divalent (Ca^2+^ and Mg^2+^) cations, and ion permeability depends on the identity of the OR, too (Butterwick et al., 2018). Under physiological conditions, ORs are localized in the dendrite of olfactory sensory neurons where they depolarize the membrane upon ligand-binding. This depolarization activates voltage-gated ion channels and changes the firing rate of the neuron (Schmidt and Benton, 2020).

An olfactory panel consisting of several ORs selected for the specific sensing of the target molecules could work effectively as a biosensor. By using the antenna of the fruit fly it was possible to distinguish between the “smell” of healthy and cancer cell lines based on their different OR activation pattern (Strauch et al., 2014). Whole antenna-based measurements however are not compatible with a sensor for the analysis of multiple samples. Several research groups conducted experiments with heterologously expressed or purified ORs linked to sensor devices, but the utilization of a complex insect receptor panel has not been realized till today. As a first step in this direction, the ligand profiles of the sensory proteins have to be characterized. However, complex information on ligand profiles, the corresponding ORs, and their genetic code is only known for a few insect species including the fruit fly (*Drosophila melanogaster*) and the malaria mosquito (*Anopheles gambiae* s.l.). The OR repertoire of both species was studied in detail *in vivo*, primarily by single sensillum recordings (Galizia et al., 2010) and by transient protein expression in *Xenopus* oocytes (Wang et al., 2010).

For our research on plant- and plant pathogen-derived VOCs (Hamow et al., 2021) *Drosophila*, due to its lifestyle, appeared to be a better starting point for the construction of biosensors than the blood-sucking malaria mosquito. Here, we report on the generation of a novel cell-based olfactory panel in which seven *Drosophila* ORs were expressed in stable human embryonic kidney (HEK) cell lines. We present the concentration-response profiles of these biosensor cell lines to a set of 66 distinct plant-related VOCs and OR-specific reference ligands. VOC-specific cell responses were determined by a high-throughput fluorescence microplate reader assay over a concentration range of three orders of magnitude. The new bio-receptor cell lines and hitherto untested plant disease-related volatiles represent original contributions to the present panel and ligand list of the examined ORs.

## 2. Materials and methods

### 2.1. Molecular cloning of OR genes into plasmid and BAC expression vectors

The OR genes OR7a (FlyBase ID FBtr0071186), OR47a (FBtr0088111), and a codon-optimized version of ORCO (FBtr0113193) were synthesized (Thermo Fisher Scientific) as double-stranded linear gene fragments. The cDNAs of the OR10a, OR13a, OR19a, OR47b, OR49b, OR67b, OR69a, OR71a, OR85b, OR98a genes were cloned from the wild-type Canton S strain of *D. melanogaster*. Total RNA was extracted from adult flies by TRIzol Reagent (Thermo Fisher Scientific) according to the manufacturer’s instructions. Total RNA (1 μg) was treated with DNase I, then reverse transcribed with oligo(dT)_12-18_ primers and SuperScript IV enzyme (Invitrogen). The OR genes were amplified with gene-specific primers containing an *Asc*I recognition sequence on both primers (except *Swa*I for OR19a) and a Kozak consensus sequence on the forward primers (Supplementary Table 1).

The above 12 OR genes linked to a wild-type internal ribosome entry site from the encephalomyocarditis virus (EMCV IRES) and the mCherry fluorescent protein gene (Shaner et al., 2004) were inserted into a modified pBlueScript KS (Agilent/Stratagene) called plasmid “L” (Zboray et al., 2015). The OR genes were inserted into this vector at the *Asc*I (*Swa*I) restriction site, while IRES-mCherry at the *Pme*I restriction site and the resulting plasmids were named as MIO(receptor name)p. The N-terminus of ORCO was fused to GCaMP6s (Chen et al., 2013) via a Gly_2_SerGly_3_SerGly linker. Subsequently, this fusion gene was cloned into a modified plasmid "L” at the *Asc*I restriction site, in which the CAG promoter (Miyazaki et al., 1989) was replaced by a Doxycycline-inducible Ptet-T6 bidirectional promoter (Loew et al., 2010) and its cognate transactivator protein (3G) gene (derived from AAVS1_Puro_Tet3G_3xFLAG_Twin_Strep, a gift from Yannick Doyon: Addgene plasmid #92099; Dalvai et al., 2015). The resulting plasmid was named as DOGGp in this work. MIO13a/47a/85b/98aDOGGp plasmids were generated by ligating *Sal*I- and *Srf*I-digested MIOp with *Bsu*36I-digested DOGGp (Figure 1A). All enzymes were purchased from Thermo Fisher Scientific (except for *Srf*I, which was from New England Biolabs). Cloning was performed in the *E. coli* DH10B strain. All plasmid sequences are available upon request.

**Figure 1.**
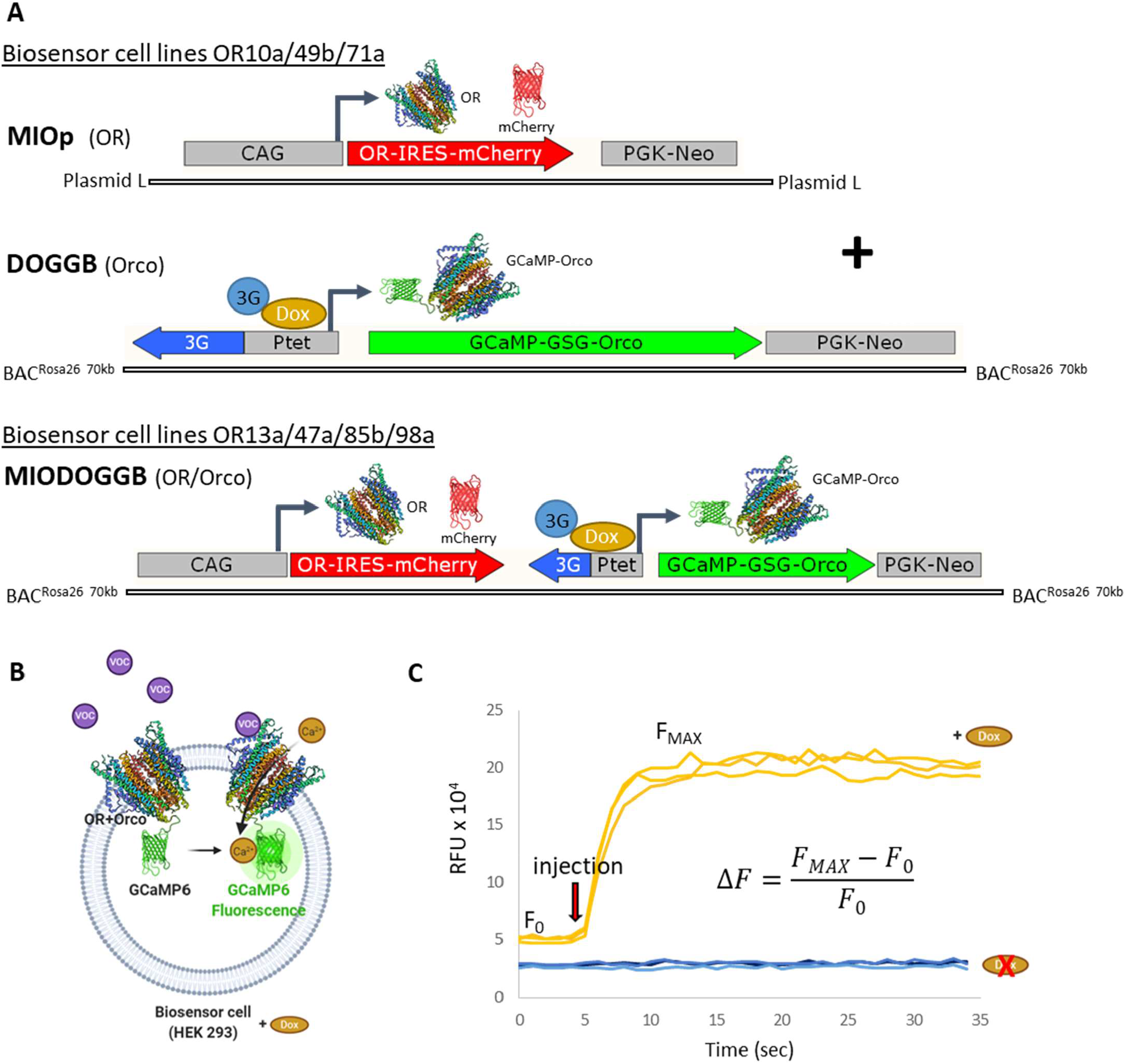
Generation and characterization of olfactory receptor-expressing biosensor cell lines. (A) DNA vector constructs for biosensor cell line production: MIOp plasmid containing the corresponding OR and mCherry, and DOGGB BAC expression vector coding for the GCaMP6-Orco fusion protein were cotransfected to generate biosensor cell lines OR10a, OR49b and OR71a (upper panel); MIODOGGB BAC expression vectors containing the corresponding OR and all other components were used to generate biosensor cell lines OR13a, OR47a, OR85b and OR98a (lower panel). The inducible Ptet bidirectional promoter only activates GCaMP6-Orco protein expression upon doxycycline (Dox) treatment. (B) Detection mechanism of the biosensor cell lines: the OR+Orco receptor complex opens upon VOC ligand binding, the resulting Ca^2+^ influx is detectable as GCaMP6-mediated fluorescence. VOC: volatile organic compound, artwork created with BioRender. (C) Fluorescence signal intensity change of the OR47a biosensor cell line 48 h after doxycycline induction (orange curves) and without doxycycline (blue curves) in response to 50 µM VUAA1. Each curve represents a technical replicate measured in separate wells of a 384-well microplate. RFU: Relative Fluorescence Unit, F_MAX_: maximal RFU value during the measurement, F_0_: average baseline fluorescence, Δ*F* was calculated according to the equation shown.

For the generation of the BAC expression vectors, a modified version of the murine *Rosa26* BAC DNA (BAC PAC Resources Children’s Hospital Oakland Research Institute clone number: RP24–85I15) was used as the vector backbone. Briefly, the original 210-kb BAC was shortened by removing approximately 70-70 kb DNA sequences from both ends of the genomic insert. First, two targeting constructs carrying 50 bp long homologous sequence regions (HR) on both ends (5’ targeting construct: HR–frt–Kanamycin–frt–HR, 3’ targeting constructs: HR– frt3–Ampicillin–frt3–HR) were generated by PCR (primers in Supplementary Table 1). The 5’ targeting construct was recombined into the targeted region replacing the original sequence by ET-cloning/Recombineering (Muyrers et al., 1999; Zhang et al., 1998) then, the cassette was excised by FLPe recombinase (Buchholz et al., 1998). Subsequently, recombination and cassette excision was repeated with the 3’ targeting construct.

DOGGp and MIO13a/47a/85b/98aDOGGp plasmids were linearized by *Sfa*AI and *Pac*I restriction enzymes to generate the linear fragments required for BAC recombination into the 2nd exon of the *Rosa26* gene by ET-cloning/Recombineering as described in detail (Zboray et al., 2015). The obtained BAC vectors were designated as DOGGB and MIO13a/47a/85b/98aDOGGB, respectively (Figure 1A).

### 2.2. Generation of biosensor cell lines

HEK293 cells (ATCC CRL-1573™) were cultured in growth medium (Dulbecco’s Modified Eagle Medium with high glucose, L-glutamine, and sodium pyruvate supplemented with 10% tetracycline-free Fetal Bovine Serum from Biosera and penicillin-streptomycin mixture). Cells were kept at 5% CO_2_ and 37°C in a humidified incubator and were passaged two-three times a week. Purified BAC and plasmid DNA were transfected into HEK293 cells with FuGene HD transfection reagent (Promega) in a 1:3 DNA:FuGene HD ratio. G418 antibiotic (Thermo Fisher Scientific) was added to the cells 48 h after transfection at 100 μg/ml concentration. For stable cell pool generation, cells were cultured for two weeks, then the antibiotic concentration was increased to 150 μg/ml and cells were cultured for one more week or until they divided vigorously without any visible sign of cell death.

Actively growing transfected cell pools were trypsinized and resuspended in 1 ml growth medium for FACS sorting. The OR13a/47a/85b/98a cell lines were sorted for mCherry fluorescence without doxycycline addition, while the OR10a/49b/71a cell lines were sorted 48 h after doxycycline addition for GFP and mCherry fluorescence with a FACSAria III cell sorter (BD Biosciences) into 96-well cell culture microplates. Clonal cell lines were cultured in growth medium supplemented with 150 μg/ml G418 and observed under an Eclipse Ti2 inverted fluorescence microscope (Nikon) when starting to grow in the 96-well microplates. Four to six actively growing mCherry positive cell clones were selected and expanded for testing their VOC response. Odorant-responsive clonal cell lines were selected and cryopreserved for subsequent experiments.

### 2.3. Calcium fluorescence assay

Twelve thousand to 18,000 biosensor cells were plated into each well of a black wall, transparent bottom 384-well microplate (Greiner) 48 h or 72 h before the measurements and were simultaneously treated with 1 μg/ml doxycycline. On the day of the experiment, the growth medium was replaced with the assay buffer (140 mM NaCl, 5 mM KCl, 2 mM CaCl_2_, 1 mM MgCl_2_, 5 mM HEPES-Na, 50 mM glucose, pH 7.3, 340 mOsm). The fluorescence was excited at 485 nm and the emission was recorded at 535 nm with a SpectraMax iD3 multimode microplate reader (Molecular Devices) mounted with a two-channel injector. Baseline fluorescence was recorded for 5 sec then VOC buffer (140 mM NaCl, 5 mM KCl, 2 mM CaCl_2_, 1 mM MgCl_2_, 5 mM HEPES-Na, 31 mM glucose, 0.1% DMSO, pH 7.3, 340 mOsm) was injected into a well and the fluorescence response was recorded for an additional 30 sec, at a rate of one read/sec. VOC solutions (four times concentrated VOCs diluted in the VOC buffer) were measured on the cells in a second round with the same protocol. Each measurement was triplicated in three wells on the same microplate, and the fluorescence intensity changes (ΔF) were calculated (Figure 1C).

### 2.4. Chemicals and chemical dilutions

All VOCs were purchased from Sigma-Aldrich (except for (5Z)-octa-1,5-dien-3-ol from Toronto Research Chemicals) in the highest available purity. VOCs were diluted either directly in the VOC dilution buffer or first in DMSO for water-insoluble compounds.

## 3. Results and discussion

### 3.1. Generation of the biosensor olfactory panel

*In vivo* measured ligand profile data derived from the DoOR database (http://neuro.unikonstanz.de/DoOR/default.html, Galizia et al., 2010; Münch and Galizia, 2016) were used to select 12 ORs, which can be appropriate to detect the VOCs relevant for our study. To assemble a novel cell-based olfactory panel, stable cell lines were successfully generated which expressed 11 different *Drosophila* ORs and the fluorescent calcium indicator protein GCaMP6 in HEK293 cells. Out of these 11 cell lines, seven were responsive to the respective OR-specific VOCs and constituted an olfactory panel in which binding of the examined odorants can be quantitated by a fluorescence signal.

Four different proteins were expressed in each of the biosensor cell lines: the respective ORs, the ORCO universal co-receptor, the mCherry fluorescent protein as a marker for OR expression, and the GCaMP6 indicator protein. To ensure stable and high expression, we used modified BAC^Rosa26^ expression vectors containing an approximately 70-kb long open chromatin region.

The expression of the OR and mCherry proteins was driven by the strong and constitutive CAG promoter (Miyazaki et al., 1989). The remaining two proteins, ORCO and GCaMP6, were expressed as a single fusion protein by linking GCaMP6 N-terminally to ORCO for membrane localization. Based on the experience of Corcoran et al. (2014) constitutive ORCO expression can lead to cell death after several weeks. In order to ensure long-term maintenance of our biosensor cell lines, the doxycycline-inducible Ptet-T6 promoter (Loew et al., 2010) was used to drive the expression of the GCaMP6-ORCO fusion protein.

Biosensor cell lines of the olfactory panel contain all of the above-mentioned genetic elements, however, in two different formats. In the OR13a, OR47a, OR85b, and OR98a cell lines the two expression units were physically integrated tandem (Figure 1A) into BAC^Rosa26^. This arrangement resulted in no baseline expression of the Ptet-T6 promoter. Based on binding experiments with the ORCO-activating artificial ligand VUAA1 (Jones et al., 2011) there was no GCaMP6-ORCO expression without doxycycline treatment whereas a strong induction was recorded 48 h after doxycycline treatment (Supplementary Figure 1A). The OR10a, OR49b, and OR71a cell lines, on the other hand, were generated by cotransfection of the same two expression units (Figure 1A). In these cell lines, the Ptet-T6 promoter had a substantial baseline activity which was further increased upon doxycycline induction (Supplementary Figure 1B). Nevertheless, this leaky promoter activity did not hinder the maintenance of these cell lines as their responsiveness was fairly stable during the 60 days monitored (Supplementary Figure 2). In the light of these results, we cannot confirm the previous observation that constitutive ORCO expression is detrimental in long-term cell culture (Corcoran et al., 2014). The cell lines may have different sensitivity to constitutive ORCO expression because the ion permeability of the channel depends on the OR identity (Butterwick et al., 2018). Alternatively, components in the culture medium may activate the ORs in some cell lines, but not in others, due to their different ligand profiles.

In four stable cell lines (OR7a, OR19a, OR69a, and OR47b) GCaMP6 and mCherry fluorescence signals were strongly visible, yet no specific ligands were identified which could activate these ORs although all these lines responded to VUAA1. Similar observations were made with heterologous expression of *A. gambiae* olfactory receptors (AgORs): 27 out of the tested 72 AgORs did not give any VOC-specific response in the *Xenopus* oocyte expression system (Wang et al., 2010).

### 3.2. The responsiveness of the biosensor cell lines, their reference ligands, and corresponding concentration-dependent OR responses

The functionality of the seven biosensor cell lines was verified by the artificial ORCO agonist VUAA1 and at least one OR-specific odorant (Figure 2). We used the GFP-linked calcium indicator GCaMP6s, which has a rise time of 100-150 msec (Chen et al., 2013), in order to follow the kinetics of odorant-binding as a fluorescence signal with a good time resolution and high-throughput (Figures 1B and 1C). Rinker et al. (2012) earlier screened a large compound library, but only one mosquito OR cell line was used in their study. Using a complex olfactory panel, we mapped here, for the first time, the concentration-response profile of a large set of defined plant- and plant disease-related odorants. The expression level of biosensor cells can show some variation between different measurements. To minimize any VOC response differences due to this variation ΔF values were normalized to average cell line-specific ΔF values in response to 50 μM VUAA1 measured on the same day and were counted as 1.

Maximum measured ΔF values on the biosensor cell lines were prominent: for VUAA1 they ranged between 374% and 541% and for reference ligands between 118% and 429%, i.e. two to ten times more than the ca. 50% previously measured in comparable experimental systems (Mitsuno et al., 2015; Termtanasombat et al., 2016).

To test the functionality of the biosensor cell lines and to identify their reference ligands, we chose two or three strong ligand candidates based on the DoOR database (http://neuro.uni-konstanz.de/DoOR/default.html; Münch and Galizia, 2016) and tested them at 100 μM final concentration. Compounds triggering the highest biosensor cell response were chosen as the reference ligand for the given OR. For four cell lines (OR49b, OR71a, OR85b, and OR98a) three odorants from our plant disease-related ligand list (see section 3.3. below) triggered stronger responses than their best ligands according to the DoOR database. In these cases, we chose these odorants as reference ligands (Figure 2B, Supplementary Table 2).

**Figure 2.**
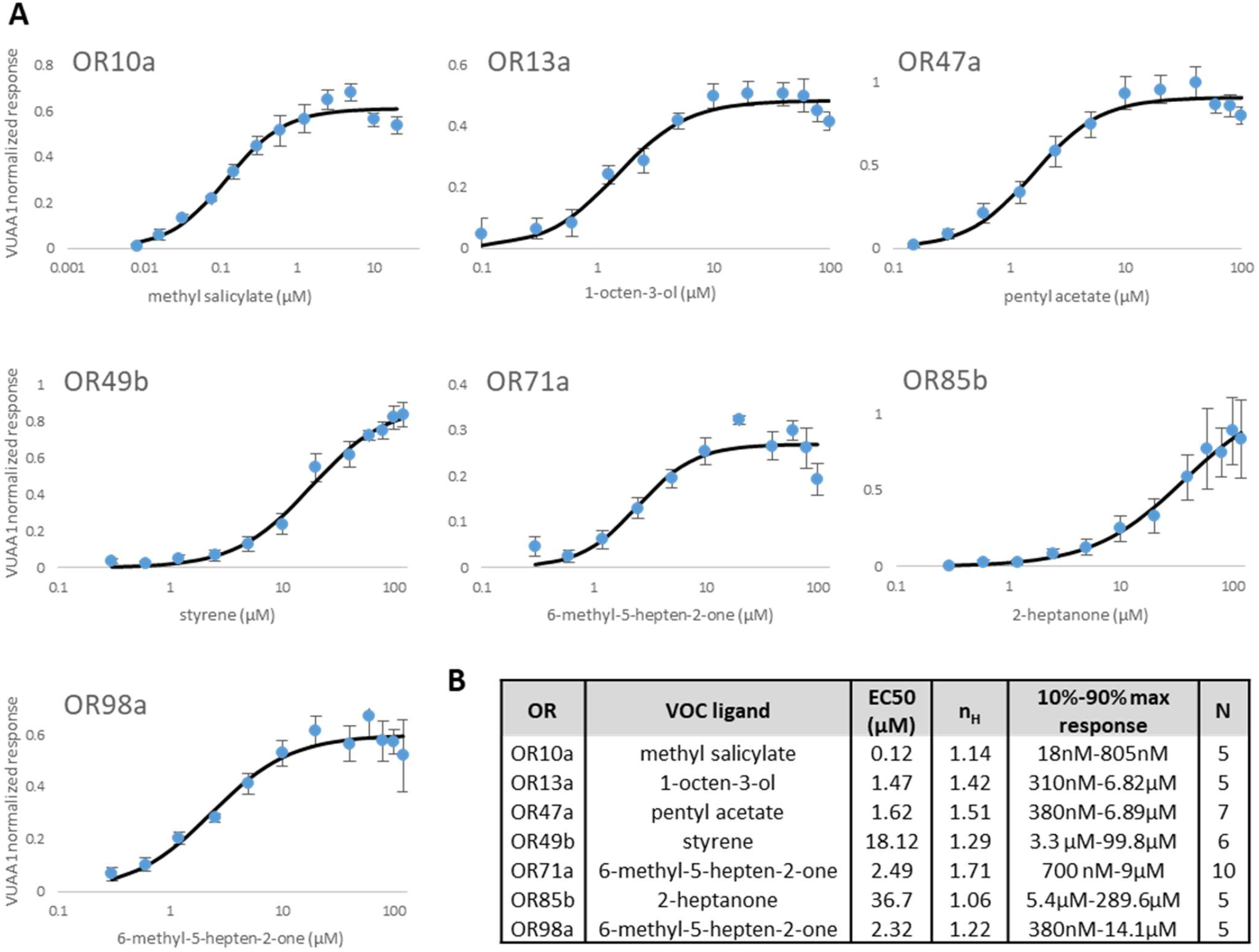
Concentration-dependent response of seven biosensor cell lines to the respective reference VOC ligands. VUAA1-normalized response was calculated by dividing VOC response with the average of VUAA1 response on the day of the measurement for the corresponding biosensor cell line. Concentration-response curves are shown as the average of N measurements of three technical replicates measured on a single day, error bars represent the standard error of the mean (SEM). EC50 values and Hill coefficients (n_H_) were calculated from the fitted Hill equation.

Out of the seven responsive biosensor cell lines reported in our study only one, OR13a was previously generated as a stable insect cell line (Sf21; Termtanasombat et al., 2016). Some other ORs were expressed only transiently either by mRNA injection into *Xenopus* oocytes or by transfection into immortalized cell lines: OR10a and OR71a (Khadka et al., 2018; Murugathas et al., 2019), OR47a (Miazzi et al., 2019; Röllecke et al., 2013; Sato et al., 2008), OR49b (Kolesov et al., 2021; Röllecke et al., 2013), and OR85b (Misawa et al., 2010; Nichols and Luetje, 2010). In contrast, all cellular biosensors generated in this study are stable cell lines that can be used for months for direct measurements. Moreover, the cell cultures can be scaled up, then cryopreserved and thawed again at any time for subsequent measurements.

We examined the concentration-dependent binding of the reference ligands on their cognate biosensor cell lines in more detail: the half maximal effective concentration (EC50) as well as Hill coefficient (n_H_) values, and the dynamic range of detection (10-90% response) were also calculated (Figure 2B). Based on these parameters a comparative evaluation of these cell lines is performed here with similar studies wherein the relevant data were available.

Concentration-dependent binding of 1-octen-3-ol in an OR13a expressing stable insect cell line resulted in an EC50 of 4.33 μM (Termtanasombat et al., 2016). Our OR13a biosensor cells were more sensitive to 1-octen-3-ol with an average EC50 of 1.47 μM (Figure 2B).

The sensitivity of the OR47a to pentyl acetate varied between different studies. In *Xenopus* oocytes, a range of 50 μM-300 μM could trigger receptor responses (Sato et al., 2008), while in a similar system the calculated EC50 was 10.7±2.0 µM (Röllecke et al., 2013). In our OR47a biosensor cell line, the average EC50 was 1.62 μM for pentyl acetate (Figure 2B), which corresponds to about seven times higher sensitivity.

For the OR49b biosensor cell line, we used styrene as a reference ligand (see section 3.3 below) but measured the EC50 for 2-methylphenol (*o*-cresol), too. With 16.52 μM of EC50 our OR49b cell line was 14-times more sensitive to this ligand than previously measured in *Xenopus* oocytes in which the calculated EC50 was 239±76 µM (Röllecke et al., 2013). In a recent study, the same OR was expressed transiently in HEK293T cells, which reacted to 2-methyl-phenol above the limit of detection in a 2 μM to 800 μM concentration range. Though the EC50 was not calculated, 80 μM concentration triggered a 38.5% response (Kolesov et al., 2021).

The OR85b receptor response to 2-heptanone was measured previously in *Xenopus* oocytes by Nichols and Luetje (2010) with an EC50 of 70±20 μM, and by Misawa et al. (2010) with an average EC50 of 45.6 μM compared to the average 36.7 μM value we obtained (Figure 2B).

### 3.3. Plant disease-related VOC measurements on the biosensor cell lines

Once responsive cell lines were selected, we tested them for sensing fungal pathogens (powdery mildew, *Botrytis cinerea, Fusarium* spp., *Pyrenophora* spp., and gray mold) of important crop plants (wheat, barley, grape, lettuce, rape, spinach, and strawberry). To this end, we had compiled a pathogen-related VOC catalog primarily based on our GC-MS analyses of infected samples (Hamow et al., 2021, and unpublished data) and also systematically retrieved from published literature data.

Together with strong ligands for our seven ORs according to the DoOR database a set of 66 compounds from our compilation were measured over a concentration range of three orders of magnitude (1 μM – 10 μM – 100 μM) on the biosensor cells (Figure 3, Supplementary Table 2). Confirming previous studies (Wang et al., 2010), the lowering of VOC concentration also decreased the number of VOCs sensed by an OR. Biosensor cells reacted to the following number of VOCs (out of 66) at 1, 10 and 100 μM of final concentrations: Or10a – 10/13/18, OR13a – 5/9/14, OR47a – 5/12/17, OR49b – 3/6/17, OR71a – 1/5/6, OR85b – 3/6/10, and OR98a – 9/8/19.

**Figure 3.**
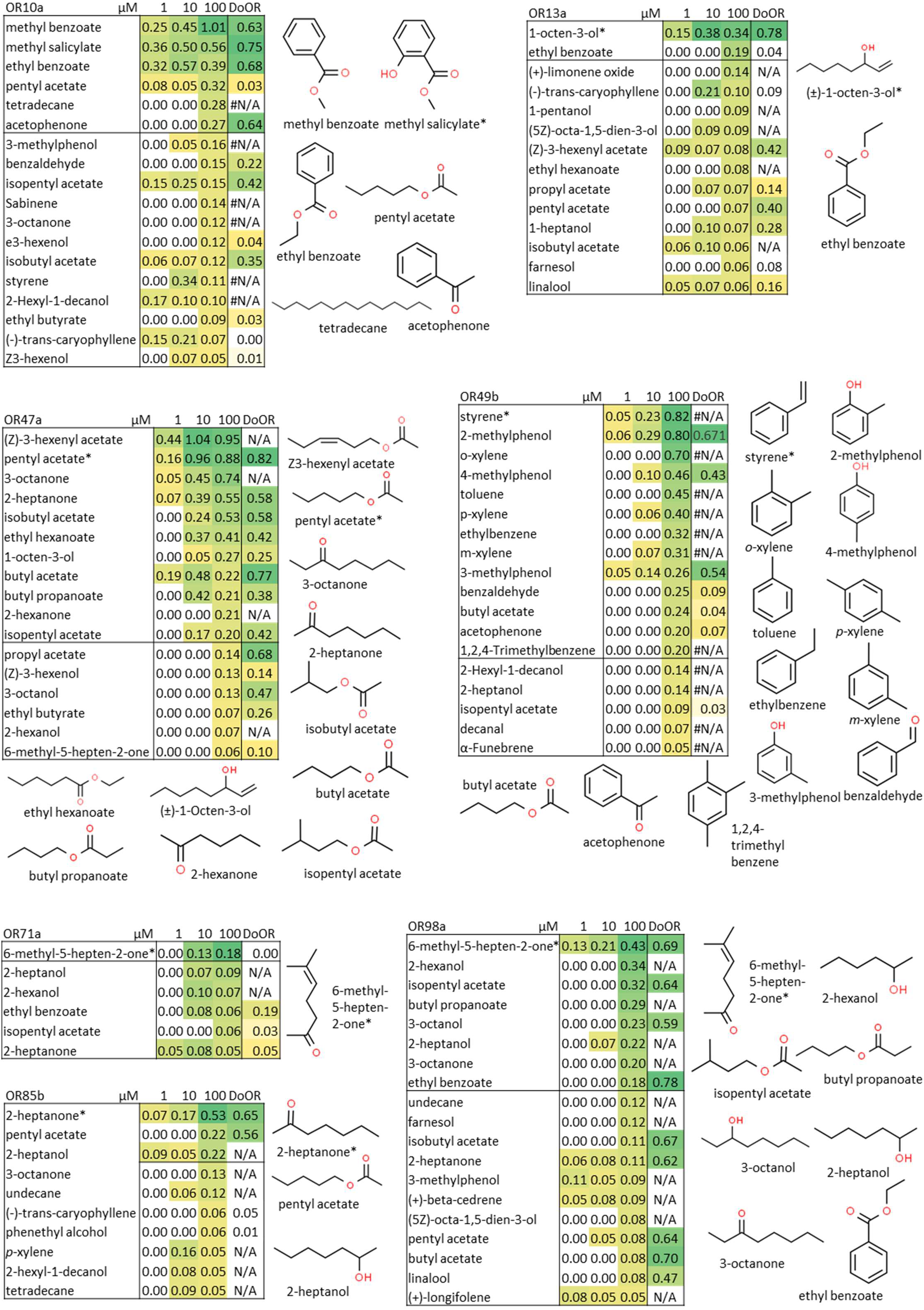
VOC ligand profile of seven biosensor cell lines. Values are averages of at least two measurements in three technical replicates. Each VOC was measured in 1, 10, and 100 µM final concentration on each biosensor cell line. VUAA1-normalized response was calculated by dividing VOC response with the average VUAA1 response on the day of the measurement for the corresponding biosensor cell line. The last (DoOR) columns contain the average response of the same olfactory receptors from the DoOR database. The chemical structures of the strong and medium ligands (at least 0.18 VUAA1-normalized response) are shown for each cell line. The reference ligands are marked with an asterisk. Heat map: white-yellow-green colors correspond to the lowest-midpoint-highest values measured for each biosensor cell line

Of all measured 1386 VOC-concentration-OR combinations we recorded 196 (14%) positive odorant responses. The majority (66.3%) of these responses were of low intensity, i.e. less than 18% of VUAA1-normalized response. Strong (at least 40% VUAA1 response) and medium (18-40% VUAA1 response) intensity reactions were in 12.2% and 21.4% of all positive cases, respectively. As the reference ligand with the lowest overall VUAA1-normalized response (6-methyl-5-hepten-2-one) elicited an 18% VUAA1-normalized response (OR71a, Figure 3), this value was taken as the threshold between low and medium/strong intensity reactions.

In 26 (5.6%) out of the 462 VOC-OR combinations tested, biosensor cell lines responded over the whole measured concentration range. High sensitivity to a ligand at the lowest concentration did not always show an overall linear relation with ligand response intensity. For example, linalool activated OR13a already at 1 μM, yet response intensity remained constant at 5-7% over the whole concentration range (Supplementary Table 2). In contrast, several odorants activated ORs only at the medium concentration, but fluorescence response was further increased at the highest concentration. As an example, isobutyl acetate activated the OR47a biosensor cell line first at 10 μM and increasing the ligand concentration to 100 μM resulted in a response intensity increase from 24% to 53% (Supplementary Table 2). Some biosensor cell lines (and their ORs) showed a well-defined affinity for structurally similar compounds. For instance, OR49b was highly specific for aromatic compounds as strong or medium responses were elicited by only these, with butyl acetate as the only exception. Figure 3 summarizes VOC ligand profiles of the biosensor cell lines and chemical structures of strong and medium ligands.

The DoOr database-derived response profiles, which are collections of *in vivo* measured datasets, served as a good starting point for the identification of candidate ORs and the prediction of their response profiles to our VOC ligands.

Twenty-two out of 66 tested VOC ligands are completely new for the biosensor cell ORs, as these are hitherto not available in the DoOR database. OR10a was the best-studied receptor for our VOC ligand set; the DoOR database contained response values for 41 VOCs, while OR71a was the least studied with only 21 previously measured VOCs. The OR10a, OR47a, and OR49b biosensor cell lines responded at 100 μM to all VOCs that were predicted as a strong ligand (at least 0.4 response) for these ORs according to the DoOR database (5/5, 8/8 and 2/2, resp. in Supplementary Table 2). Similarly, a high matching rate (9/15, 60%) was also found for the OR98a biosensor cell line. However, for two cell lines, OR13a and OR85b, the respective success rates were only 33% (2/6) and 20% (2/10), and the ligand profile of the OR71a biosensor cell line did not match at all the DoOR-based predictions. These observations indicate the importance of testing the ligand profile and sensitivity in a different expression system.

Recently, powdery mildew-specific volatile biomarkers have been quantified in the headspace of healthy and inoculated wheat plants (Hamow et al., 2021). VOCs from the headspace of diseased plants were collected on a Porapak Q sorbent and eluted with DMSO for biosensor response profiling. The VOC biomarker concentrations in these DMSO samples were highly variable as determined by GC-MS (Table 1). Three VOCs, 1-octen-3-ol, (5Z)-octa-1,5-dien-3-ol, and 3-octanone were in the detectable range for samples from heavily infected plants, while the concentration of 1-heptanol was below the detection limit of the OR13a biosensor cell line.

**Table 1.**
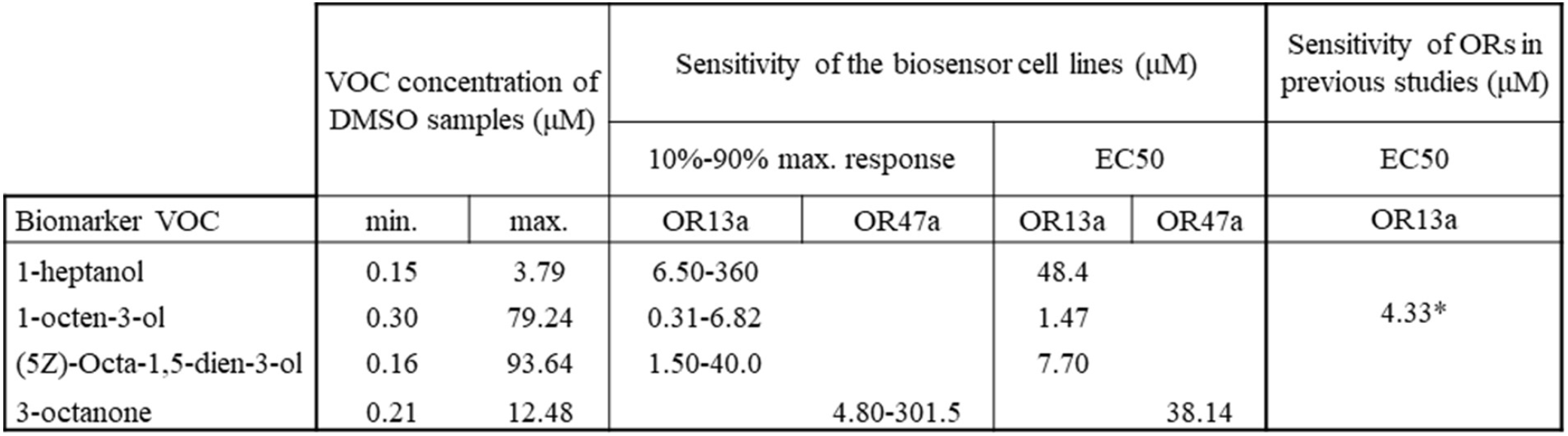
Concentration range and detectability of major wheat powdery mildew biomarker VOCs by two biosensor cell lines. *Sf21 (insect) stable cells (Termtanasombat et al., 2016)

The current time requirement for the profiling of an unknown sample was only 13 min with our biosensor panel when measured in triplicates (three wells/sample/biosensor cell line). On a whole 384-well microplate it was possible to measure 16 samples within ca. 3.5 h.

We analyzed three healthy wheat-derived and seven powdery mildew-infected plant-derived samples with a microplate reader assay. Due to our protocol, the samples contained DMSO and had to be diluted at least 200-fold before the measurements. However, with this degree of dilution, no significant differences were observed between healthy and infected plant samples. Sample collection and measurement protocol for such plant-derived material require further optimization for the successful distinction between healthy and infected samples.

The OR elements of the cellular biosensors introduced in our study can also be extracted from the cellular system and coupled to other types of signal-transducing elements. For instance, systems like the ones developed by Murugathas et al. (2019) or Yamada et al. (2021) may be utilized for our biosensors, too, and such coupling steps could further improve sensitivity for the ligands explored here.

## 4. Conclusions

We generated responsive biosensor cell lines which stably express seven functional insect ORs together with GCaMP6, a fluorescent calcium indicator protein. These cell lines were produced by transfection with a novel BAC expression vector carrying an open chromatin region to ensure higher and more stable expression levels in a long term. The biosensor cell lines detected odorant molecules from the ppb concentration in the liquid phase. The response was quick with the cells reaching maximum fluorescence intensity change within 10 sec. Peaks of fluorescence intensity change were as high as 541% for the ORCO agonist VUAA1 and 429% for reference ligands, which is outstanding compared to published data of ca. 50%. The concentration-response profile of the biosensor cell lines was mapped for 66 plant- and plant pathogen-derived VOCs over a concentration range of three orders of magnitude. We recorded ligand-specific fluorescence responses in 14% of all 1386 measured combinations of VOC concentrations and biosensor cell lines. Three out of the four major volatile biomarkers from powdery mildew-infected wheat were detectable in undiluted samples, but not in 200-fold diluted ones, by the biosensor cells. The ORs expressed by the biosensor cell lines can be purified for coupling to other signal-transducing devices in order to achieve even higher sensitivity.

## Acknowledgments

This work was supported by the Economic Development and Innovation Operational Programme (grant No. GINOP-2.3.2-15-2016-00051). Cell sorting was conducted by György Várady at the Laboratory of Flow Cytometry of the ELKH Research Centre for Natural Sciences (Budapest). We thank Bettina Menich and Zainab Quddoos for technical assistance in cell maintenance and microplate reader measurements as well as Orsolya Németh and Anikó Keszőcze for help in OR cloning.

