## Supplementary material for "High-throughput ligand profile characterization in novel biosensor cell lines expressing seven heterologous olfactory receptors for the detection of volatile plant biomarkers"

### Supplementary Table 1.

*Primer list for cDNA cloning of olfactory receptor (Or) genes*

| Name | Sequence (5’-3’) |
|---|---|
| Or10a for | TTTTTGGCGCGCC**GCCACCATG**TCCGAGTGGTTACGC |
| Or10a rev | AAAAAGGCGCGCCTTACTGAAAGGACTTAACCAGC |
| Or13a for | TTTTTGGCGCGCC**GCCACCATG**TTCTATTCGTATCCCTAC |
| Or13a rev | AAAAAGGCGCGCCTTAATCTAGTTTCTTTTCGTCG |
| Or19a for | TTTTTATTTAAAT**GCCACCATG**GACATATCGAAGGTGG |
| Or19a rev | AAAAAATTTAAATCTATTCAAGGGACGTTTTACG |
| Or47b for | TTTTTGGCGCGCC**GCCACCATG**AACGACTCGGGTTATC |
| Or47b for | AAAAAGGCGCGCCCTACATCGATTCTTGCATCAG |
| Or49b for | TTTTTGGCGCGCC**GCCACCATG**TTTGAAGACATTCAGC |
| Or49b rev | AAAAAGGCGCGCCTCATCCGTAGACTCGCTT |
| Or67b for | TTTTTGGCGCGCC**GCCACCATG**CAGGACCAACTGGATC |
| Or67b rev | AAAAAGGCGCGCCCTATTTGTTGTTCATGTTGCG |
| Or69a for | TTTTTGGCGCGCC**GCCACCATG**CAGTTGCACGACCATATG |
| Or69a rev | AAAAAGGCGCGCCTTATTTAAGGGACCGCACAC |
| Or71a for | TTTTTGGCGCGCC**GCCACCATG**GACTACGATCGAATTC |
| Or71a rev | AAAAAGGCGCGCCCTATTGGTTCATGTTGAGC |
| Or85b for | TTTTTGGCGCGCC**GCCACCATG**GAGAAGCTAATGAAGTAC |
| Or85b rev | AAAAAGGCGCGCCCTATTGGGTATACATTGTGC |
| Or98a for | TTTTTGGCGCGCC**GCCACCATG**TTGTTCAACTATCTGCG |
| Or98a rev | AAAAAGGCGCGCCTCAGTTCTTTGTCAATCTGTC |

Restriction enzyme recognition sites (*Swa*I for OR19a, *Asc*I for all other receptors) underlined, Kozak consensus sequences in **bold**.

*Primer list for the generation of the 70 kb Rosa26 BAC targeting constructs*

| Name | Sequence (5’-3’) |
|---|---|
| BAC $^{\text{Rosa26 70kb}}$ 5′ for | GAATTTTCTATATTATGAATGTCTCTGTAATAAATAAATCAATTCTTCAAGCGAAG TTCCTATTCTCTAGAAAG |
| BAC $^{\text{Rosa26 70kb}}$ 5′ rev | GATAAATACGTACTTCAAGTTTAAGAGTGAGAGAAACTTCAAGGCAGTTCAGAA GTTCCTATACTTTCTAGAG |
| BAC $^{\text{Rosa26 70kb}}$ 3′ for | TGAACCATCTTGCTAACTCACAGTGGTATTTTCTCAGGATACTTCTGTTTGAAGTT CCTATTCTTCAAATAG |
| BAC $^{\text{Rosa26 70kb}}$ 3′ rev | TGCCTCAGGATGTCTTGATTTTGGTTATCTGGGATTAAATCTACTAAGTATGGAA GTTCCTATACTATTTGAAG |

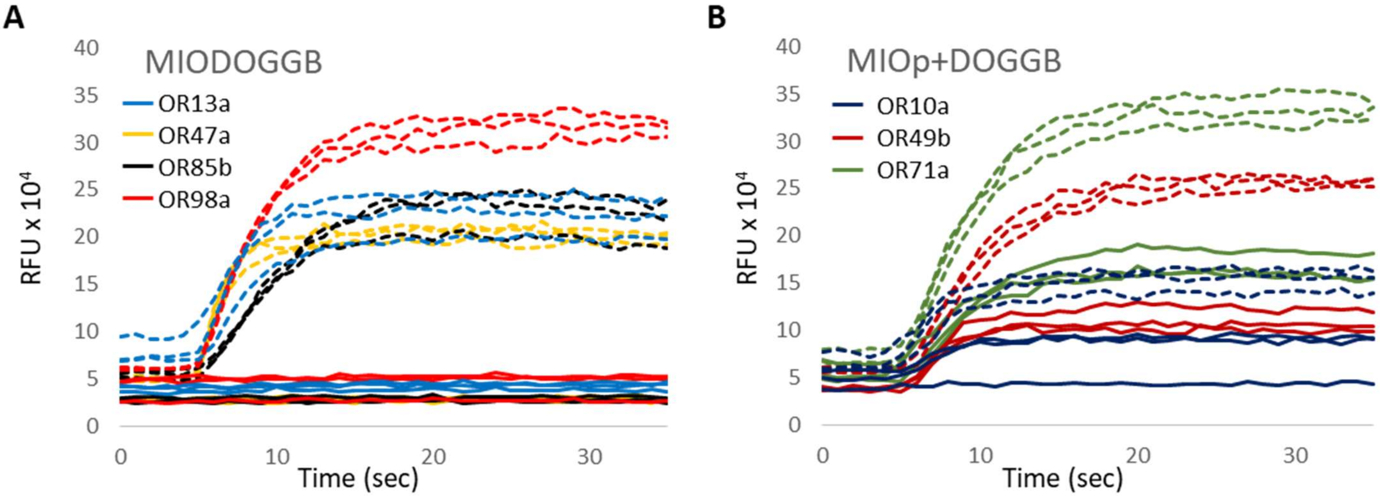


**Supplementary Figure 1. Inducible GCaMP6-Orco expression of seven biosensor cell lines.** (A) Expression of the four biosensor cell lines generated with the MIODOGGB expression vectors (Figure 1A). (B) Expression of the three biosensor cell lines generated by cotransfection of the MIOp plasmids containing the corresponding OR and the DOGGB expression vectors containing the GCaMP6-Orco fusion protein (Figure 1A). In all cases, 50 µM VUAA1 was injected into the wells and the fluorescence intensity change was recorded in three replicates. Dashed curves: GCaMP6-Orco expression induced in biosensor cells by doxycycline for 48 h, solid curves: no doxycycline added to the cells (control).

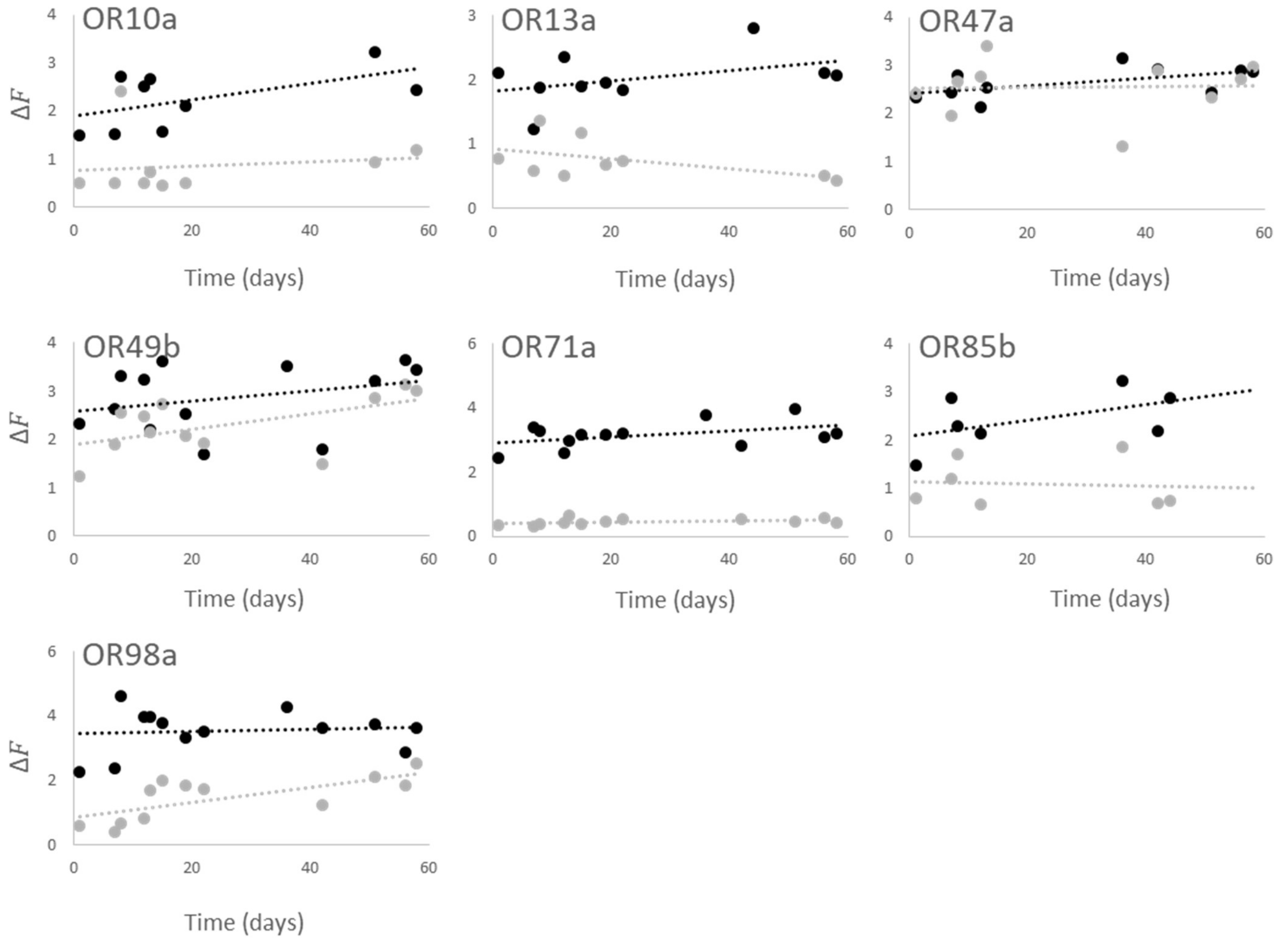


**Supplementary Figure 2. Ligand response stability of seven biosensor cell lines.** Fluorescence intensity change ($\Delta F$) of the biosensor cell lines in response to their reference ligands (light gray dots): OR10a: 10 µM methyl salicylate, OR13a: 10 µM 1-octen-3-ol, OR47a: 10 µM pentyl acetate, OR49b: 100 µM styrene, OR71a: 100 µM 6-methyl-5-hepten-2-one, OR85b: 100 µM 2-heptanone, OR98a: 100 µM 6-methyl-5-hepten-2-one, and to VUAA1 (black dots) during 60 days of culture.
